# Inside and outside of virus-like particles HBc and HBc/4M2e: a comprehensive study of the structure

**DOI:** 10.1101/2022.07.04.498777

**Authors:** V.V. Egorov, A.V. Shvetsov, E.B. Pichkur, A.A. Shaldzhyan, Ya.A. Zabrodskaya, D.S. Vinogradova, P.A. Nekrasov, A.N. Gorshkov, Yu.P. Garmay, A.A. Kovaleva, L.A. Stepanova, L. M. Tsybalova, T.A. Shtam, A.G. Myasnikov, A.L. Konevega

## Abstract

The structure of the core particles of the hepatitis B virus core antigen (HBc) with the insertion of four external domains of the influenza A M2 protein (HBc/4M2e) have been studied using a combination of molecular modeling and cryogenic electron microscopy (cryo-EM). It has also been shown that synthesis of particles occurs inside bacterial cells, but despite the big inner volume of the core shell particle, purified HBc/4M2e do not contain detectable amounts of bacterial proteins. It has been shown that a fragment of the M2e is prone to the formation of amyloid-like fibrils. However, as a part of HBc immunodominant loop M2e domains as repeats, does not exhibit a tendency to aggregation. Full-atom HBc-4M2E model with 3A resolution was obtained by molecular modeling methods based on cryo-EM data.

---

Virus-like particles (VLPs) are widely used in vaccine development [1]. The most commonly used vaccines are based on the HBc antigen of the hepatitis B virus [2]. Virus-like HBcore particles are a complex of monomers (with a molecular weight of 21 kDa), into the so-called “antigenic loop” of which the target antigen is inserted by genetic engineering methods. Such monomers are capable of self-assembly into spherical particles of two types, T3 symmetry, 180 monomers and T4 symmetry, 240 monomers [3].

For the use of HBc VLPs in vaccines, it is necessary that: (i) insertion into the “antigenic loop” was localized outside the VLP; (ii) VLPs did not show a tendency to oligomerization or aggregation; (iii) VLPs do not contain any contaminant proteins inside the shell. In this study, we tested the aforementioned properties of virus-like HBc particles with four tandem inserts in an antigenic loop constructed on the basis of the external domain of the M2 protein of the influenza A virus (HBc/4M2e). Transmission electronic and atomic force microscopy used in the study of VLPs do not have sufficient resolution to determine the position and conformation of the insert in the “antigenic loop”. In this work, to determine the position of the 4M2e insert in HBc virus-like particles, we used cryo-electron microscopy. We study the aggregation properties of the peptide corresponding to a fragment of an antigenic insert (external domain of the influenza A virus M2, M2e) and the aggregation properties of virus-like particles with four M2e-based sequence tandem inserts in the antigenic loop. Aggregation properties was studied using dynamic light scattering, and particle stability was examined with differential scanning fluorimetry. The protein composition of HBc/4M2e was characterized using mass-spectroscopy.

## MATERIALS AND METHODS

### Peptide G37

Peptide G37 (SLLTEVETPIRNEWGCRCNDSSD, 2621.15 Da), was synthesized at OOO NPF VERTA, purity more than 80%. Peptide was dissolved in phosphate buffered saline (PBS) buffer to the 1 mg/ml concentration and then incubated at room temperature for 1 hour.

### Expression and purification of HBc and HBc/4M2e particles

HBc particles have been obtained as described in [3]. Briefly, *E*.*coli* cells were sonicated, then a series of centrifugations, precipitation with ammonium sulfate, followed by purification using ion exchange chromatography and gel filtration were carried out.

### Hydrodynamic radius and zeta-potential

The measurement of the hydrodynamic radius or zeta-potential of the HBc or HBc/4M2e particles and size of G37 peptide oligomers was carried out by the dynamic light scattering using a Malvern Zeta-sizer nano spectrometer (Malvern Instruments) according to the manufacturer’s instructions.

### Atomic Force Microscopy

The G37 peptide samples were diluted 10 times with water, after which 10 μL of solution were applied onto a freshly cleaved mica substrate. After 1 minute of incubation, the samples were washed three times with 500 μL of water to remove salts and unbound components, and then dried in air. Images (topography of the sample surface) were obtained in the semi-contact mode on an atomic force microscope “NT-MDT” (NT-MDT, Russia), with NSG03 probe. Image processing was performed using “Gwy ddion” software [4].

### Transmissive electronic microscopy

At the first stage, to make the surface of the support grids for electron microscopy covered with a film of amorphous carbon (01843-F / 01844-F, Ted Pella) hydrophilic, the grids were glow-treated using a PELCO easiGlow device under the following conditions: sample processing time - 30 sec, current strength - 0.25 mA, residual pressure in the chamber - 0.26 mbar. At the next stage, 3 μL of a solution of the object of study was applied to the treated grids, which was kept for 1 min and contrasted for 1 min using a 1% solution of uranyl acetate. The finished meshes were kept for at least 15 min until complete drying before examination in a Titan 80-300 transmission electronic microscope.

### Transmission electron microscopy of bacterial sections

The bacterial suspension was precipitated by centrifugation, prefixed with 2.5% glutaraldehyde in PBS, postfixed with 1% OsO4 solution, dehydrated in a series of ethanol solutions of increasing concentration, impregnated with acetone and embedded in Epon epoxy resin. Next, ultrathin sections of these blocks were obtained, the grids with sections were contrasted with an alcohol solution of uranyl acetate, an aqueous solution of lead citrate, dried and examined in JEOL JEM 1100 microscope.

### Differential Scanning Fluorimetry

The conformational stability of HBc/4M2e on the thermal denaturation analysis was performed by differential scanning fluorimetry without additional fluorescent labels by smoothly heating the sample in a scanning fluorometer (Prometheus NT.48, NanoTemper Technologies GmbH, Germany). HBc/4M2e samples in PBS buffer solution (at concentrations of 3 mg/ml) were collected in glass capillaries (Standard Capillaries Prometheus NT.48, NanoTemper), capillaries were placed on the platform and measurements were taken in the temperature range from 15°C to 95°C in increments of 1°C /min at a laser intensity of 50%. The fluorescence intensity was recorded at 330 nm and 350 nm, and the ratio of the fluorescence intensities and the first derivative was analyzed.

### Tryptic hydrolysis in solution

Recombinant proteins HBc and HBc/4M2e were diluted with water to a concentration of 200 μg / ml, mixed 1 to 1 (v / v) with 10 μg / ml trypsin (Sequencing Grade Modified Trypsin, Promega) in 50 mM ammonium bicarbonate and incubated for 2 hours at 60 °C for enzymatic hydrolysis. The reaction was stopped by adding 2% trifluoroacetic acid (TFA) 1 to 1 (v / v).

### Mass spectrometry

Samples were applied to a target in a DHB matrix (20 mg/ml 2,5- dihydroxybenzoic acid, 30% acetonitrile, 0.5% TFA): 0.75 μl of the sample was applied to 0.75 μl of the matrix, and dried at room temperature. Mass spectra were obtained in positive ions registration mode using MALDI-TOF mass spectrometer “ultrafleXtreme”, Bruker under the control of the program “flexControl”, Bruker. For each spectrum, 3000 Nd laser pulses were summed up. The spectra were processed using the flexAnalysis software, Bruker. Proteins were identified using the Biotools software (Bruker) and Mascot (http://www.matrixscience.com).

### SDS-PAGE and staining with Coomassie colloidal solution

The initial concentration of recombinant proteins HBc and HBc/4M2e was 1.7 mg/ml. The polyacrylamide gel electrophoresis (PAGE) was carried out under denaturing conditions in the presence of sodium dodecyl sulfate (SDS) and beta-mercaptoethanol [5]. Samples were mixed with 4x Laemmli’s buffer (final concentrations: 25 mM Tris-HCl pH 6.8, 100 mM beta-mercaptoethanol, 1% SDS, 0.05% bromophenol blue, 5% glycerol) and denatured at 70 ° C for 20 minutes. 2 μl of recombinant proteins and 1 μl of a molecular weight marker (Precision Plus Protein Kaleidoscope Prestained Protein Standards, BioRad) were applied to the PAGE lane (10 wells, Any kD Mini-PROTEAN TGX Protein Gels, BioRad). The cathodic and anodic buffers were 25 mM Tris, 250 mM glycine, 0.1% SDS. EF parameters: 25mA, 220V, 50 minutes. Gels were stained with Coomassie colloidal solution (10% phosphoric acid, 10% ammonium sulfate, 0.12% Coomassie Blue, 20% methanol) [6] overnight, washed with water.

### Cryo-electron microscopy

The study of the HBc and HBc/4M2e samples by cryo-electron microscopy was carried out at the RC PEM KK NBICS using a Titan Krios 60-300 cryo-electron microscope equipped with a Falcon II high-performance electron detector using EPU (FEI) software. Supporting grids for electron microscopy with periodic holes in an amorphous carbon film (Quantifoil R1.2 / 1.3, Quantifoil) were processed in a glow discharge using a PELCO easiGlow device under standard conditions: sample processing time - 30 sec, current - 0.25 mA, residual chamber pressure - 0.26 mbar. At the first stage, samples of hepatitis virus (HBc) core particles were vitrified in the chamber of the Vitrobot Mark IV installation with the following main parameters: blot force - 0 units, blot time - from 2.5 to 3.5 sec, the temperature in the chamber is 4.5 degrees Celsius, the humidity in the chamber is 95-100%. Experimental images were obtained with the parameters given in Table 1.

**Table 1.** Reliable identified protein using mass spectrometry in HBc/4M2e sample.

| Band | Protein | Organism | Band % | M, kDa | Score (threshold) |
| --- | --- | --- | --- | --- | --- |
| #1 | bifunctional acetaldehyde-CoA/alcohol dehydrogenase; | E.coli | 3.6 | 100 | 181 (92) |
|  | pyruvate dehydrogenase (acetyl-transferring) |  |  |  | 193 (92) |
| #2 | 60 kDa chaperonin | E.coli | 0.9 | 86 | 213 (90) |
| #3 | HBc/4M2e | recombinant protein | 19.4 | 72 | 134 (92) |
| #4 | tryptophanase | E.coli | 3.3 | 48 | 147 (92) |
|  | dihydrolipoamide succinyltransferase |  |  |  | 136 (92) |
| #5 | HBc/4M2e | recombinant protein | 72.8 | 38 | 116 (92) |

Note: if Score exceed the threshold, the identification considered reliable ( $p < 0.05$ )

**Table 1. Parameters for obtaining experimental data for the subsequent determination of the structure of the object of interest using the method of analysis of projections of single particles using a cryo-electron microscope Titan Krios 60-300**

Accelerating voltage, kV 300

Nominal magnification 78000x

First condenser aperture, μm 2000

Second condenser aperture, μm 100

Objective aperture, μm 100

Exposure time, s 2.0

Defocus range, μm [-1.2: -3.0]

### Cryo-electron microscopy data processing

Two density maps were obtained for hepatitis B virus particles with an insert with two different topologies (T3 and T4) with a resolution of about 3A. Each particle was inscribed with an asymmetric unit with the appropriate topology taken from the pdb databank (for T3 - 7EOY for T4 - 7OD4) using chimeraX software, then the structure was refined using a phenix (using the rigid body fitting). For the fit structures with an insert, the fragments of the 25A inserts were modeled and then re-fit using the Phenix. All residues that did not fit into the electron density were removed.

## RESULTS AND DISCUSSION

The sequence design of the HBc/4M2e virus-like particles implied the insertion of four tandemly repeated fragments of the influenza A M2 protein outer domain (M2e) in the immunodominant loop [3], [7]. The virus-like particles suspension stability depends both on the structure and charge of the HBc particle itself and on the structure and properties of the target immunogen inserted into HBc immunodominant loop. To assess the aggregation potential of four M2e fragments in HBc/4M2e fusion protein, first we studied the aggregation properties of the peptide corresponding to the M2e antigen primary structure (G37).

A dynamic light scattering study of the solution containing G37 peptide oligomers showed that the dissolved peptide after 1 hour of incubation at room temperature contains high molecular weight aggregates with a multimodal hydrodynamic radius distribution in excess of 200 nm (Figure 1A). The multimodality of the distribution did not allow to reliably determine this parameter, so the sample was subjected to atomic force microscopy. The results of atomic force microscopy are presented in Figure 1B.

**Figure 1.**
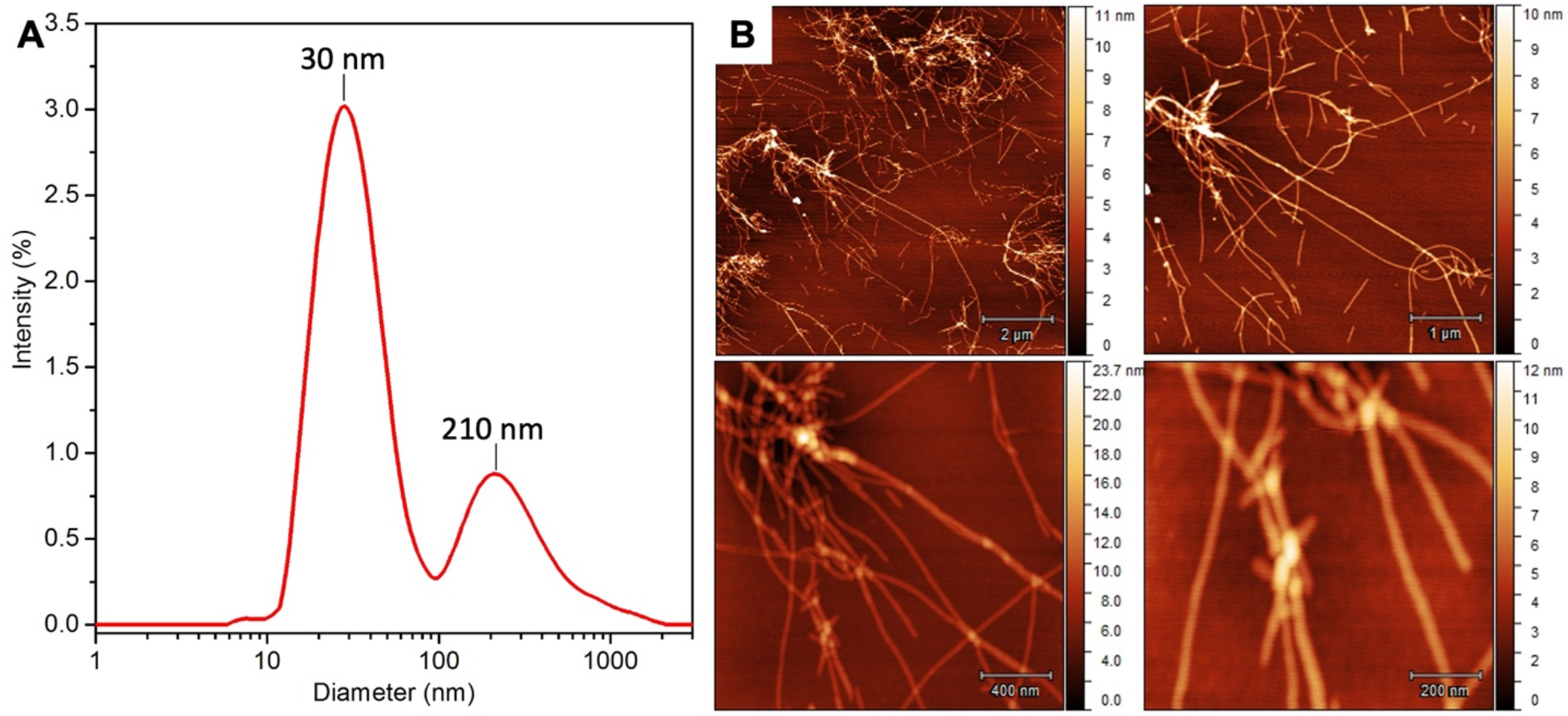
(**A**) G37 aggregate hydrodynamic ratio distribution measured with dynamic light scattering. (**B**) Atomic force microscopy of aggregates contained in a solution (1 mg/ml) of G37peptide. Above - bundles of thin short fibrils, below - long fibrils.

Since the detected fibrils were morphologically similar to amyloid-like ones (see, for example, [8]), we checked the fibrils obtained for the properties characteristic for amyloid-like ones - namely, the ability to shift the absorption spectrum of the Congo Red dye into the long-wavelength region [9] and increase the fluorescence intensity of Thioflavin T [10]. Figure 2 shows the absorption spectrum of the Congo red dye (A) and Thioflavin T (B) fluorescence spectrum in the presence and absence of aggregates of the G37 peptide. An additional peak at 550 nm in the presence of the peptide in Congo Red spectrum and more than 30x increase in ThT fluorescence intensity indicate the amyloid-like nature of the G37 peptide aggregates. Thus, it was shown that the M2e fragment of the insert in the immunodominant loop is prone to the formation of amyloid-like fibrils.

**Figure 2.**
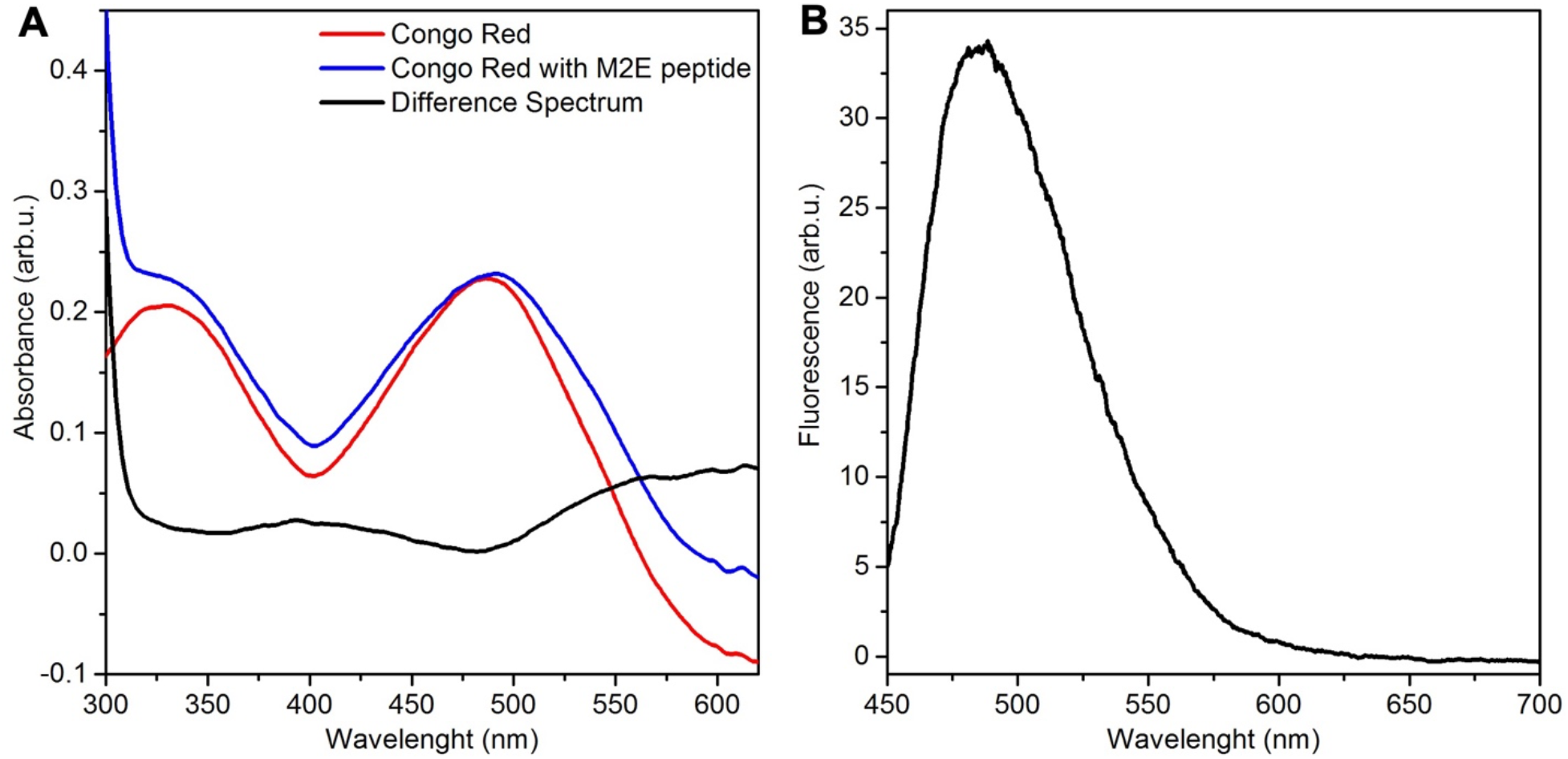
(**A**) Absorption spectrum of Congo red dye in the presence (red curve) and absence (black curve) of the G37 peptide, and difference spectrum (green curve). An additional peak at 550 nm (on the difference spectrum) in the presence of the peptide indicates the amyloid-like nature of the aggregates. (**B**) ThT spectrum in the presence of G37 fibrils, spectrum of pure ThT was used as a baseline.

Despite the amyloidogenicity of the M2e insert, HBc/4M2e particles showed stability in solution. Earlier, a number of researchers have shown that HBc particles themselves in PBS buffer have a zeta potential about -15 mV, which corresponds to their existence in the form of a metastable colloidal solution (link about zeta) and is consistent with the data on the HBc zeta potential obtained by us (−10 mV). Measurement of HBc/4M2e zeta potential showed value (− 32.1±6.3 mV) corresponding to a stable suspension. The immunodominant loop M2e insertion appears to have stabilized in VLPs.

The integrity of HBc/4M2e particles was monitored using differential scanning fluorimetry and transmissive electronic microscopy (Figure 3). By analyzing TEM images, it was found that the lifetime of HBc/4M2e particles with unchanged morphology is no more than a month. After a month of storage of the samples at +4 degrees, electron microscopic images show only particles with altered morphology, namely, with the broken integrity of the spherical shell (Figure 3A, B). The spectra of intrinsic fluorescence obtained on suspensions of such particles by the method of differential scanning fluorimetry indicate the accompanying process of violation of the integrity of changes in the polarity of the microenvironment of aromatic residues (Figure 3C, D).

**Figure 3.**
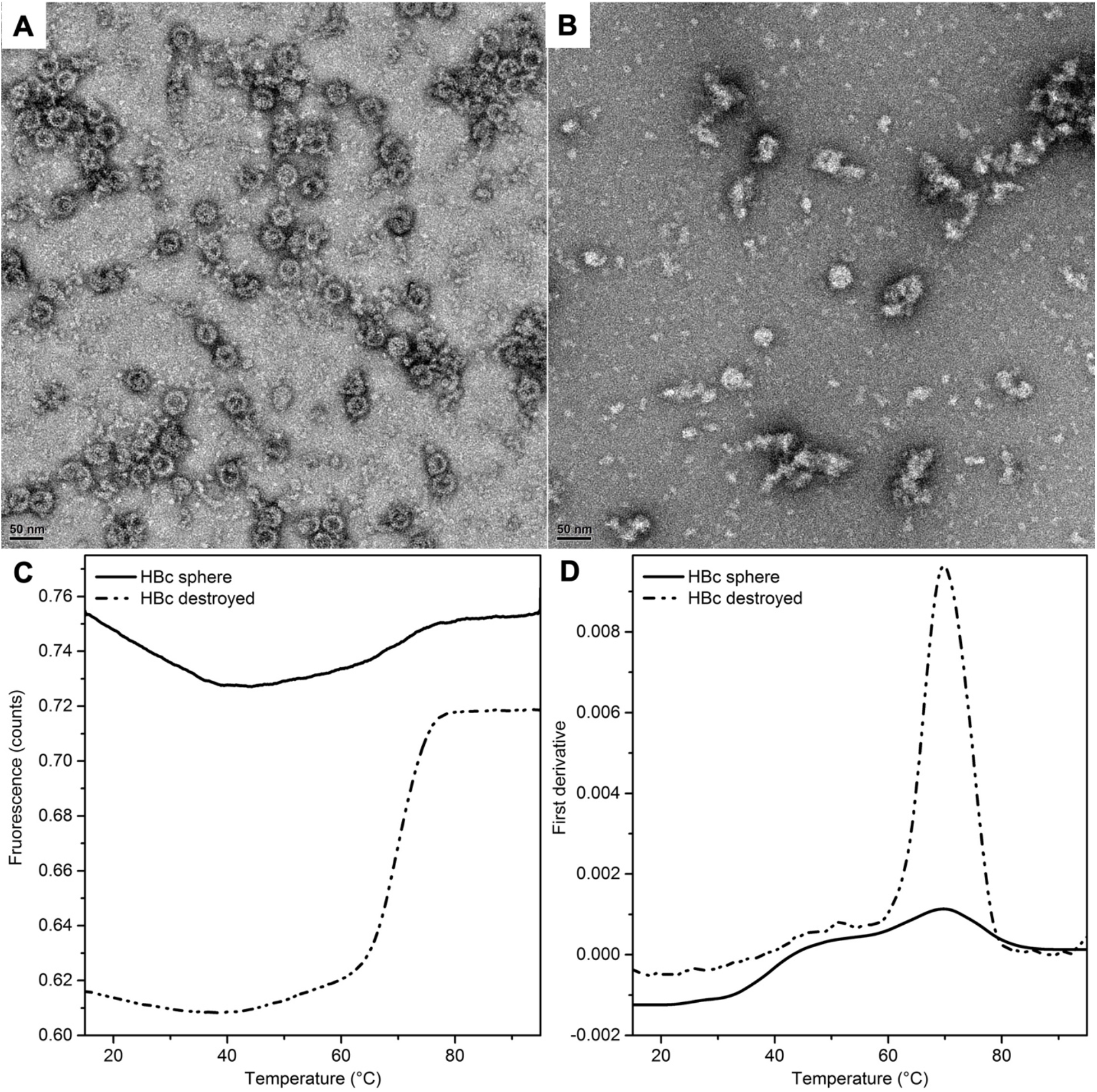
(**A, B**) Electron microscopic images of HBc/4M2e particles before and after their storage in a PBS solution at +4 ºC. (**C, D**) The results of differential scanning fluorimetry of samples of native and destroyed HBc/4M2e - the dependence of the 330/350 nm fluorescence intensity ratio on temperature (left) and the first derivative of this dependence (right). The spectrum of native particles is shown by a solid line, the spectrum of denatured ones is shown by a dotted line.

To determine the position and structure of the antigen-presenting loop the unbroken HBc/4M2e particles were subjected to cryo-electron microscopy followed by analysis of single particles due to the scheme represented in Figure 4.

**Figure 4.**
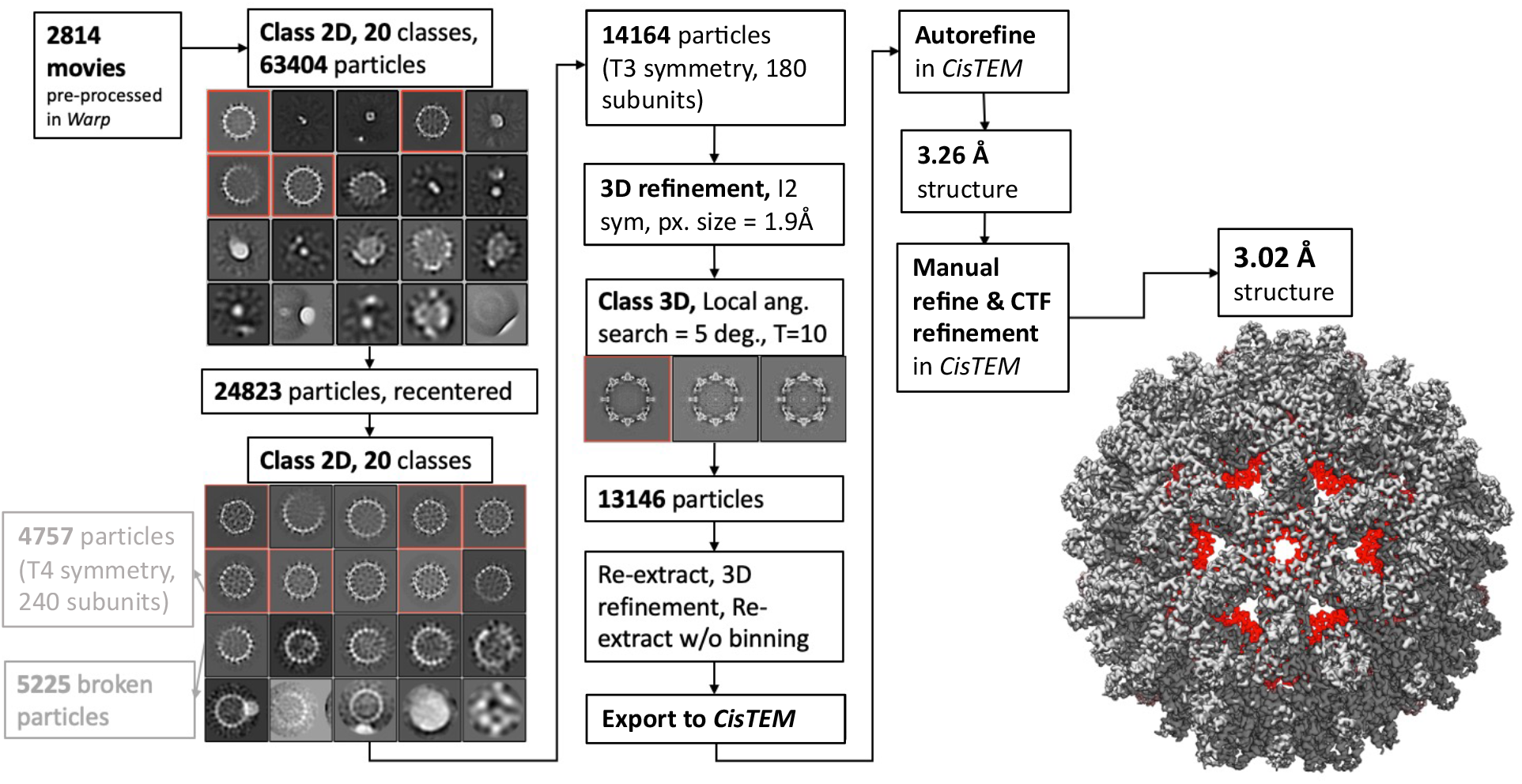
Cryo-electron microscopy of HBc/4M2e particles. Scheme of Cryo-EM data processing.

Two electron density maps were obtained for HBc/4M2e particles with two different topologies (T3 symmetry, 180 monomers, and T4 symmetry, 240 monomers per particle) with a resolution of about 3A (Figure 4). The images obtained allowed us to determine the structure of the HBc particle and the location of the beginning and the end of the 4M2e antigenic loop for HBc/4M2e particles with T3 and T4 symmetry (Fig. 5A, B). At the same time, the position of the antigenic loop structure could not be localized, apparently due to its high mobility. Similar problems were encountered by other researchers studying the loop insertion structure [11] It was possible to say unambiguously that there was no violation of the structure of the core particle, despite the tendency of the sequences contained in the antigenic loop to form beta-structured amyloid-like oligomers.

**Figure 5.**
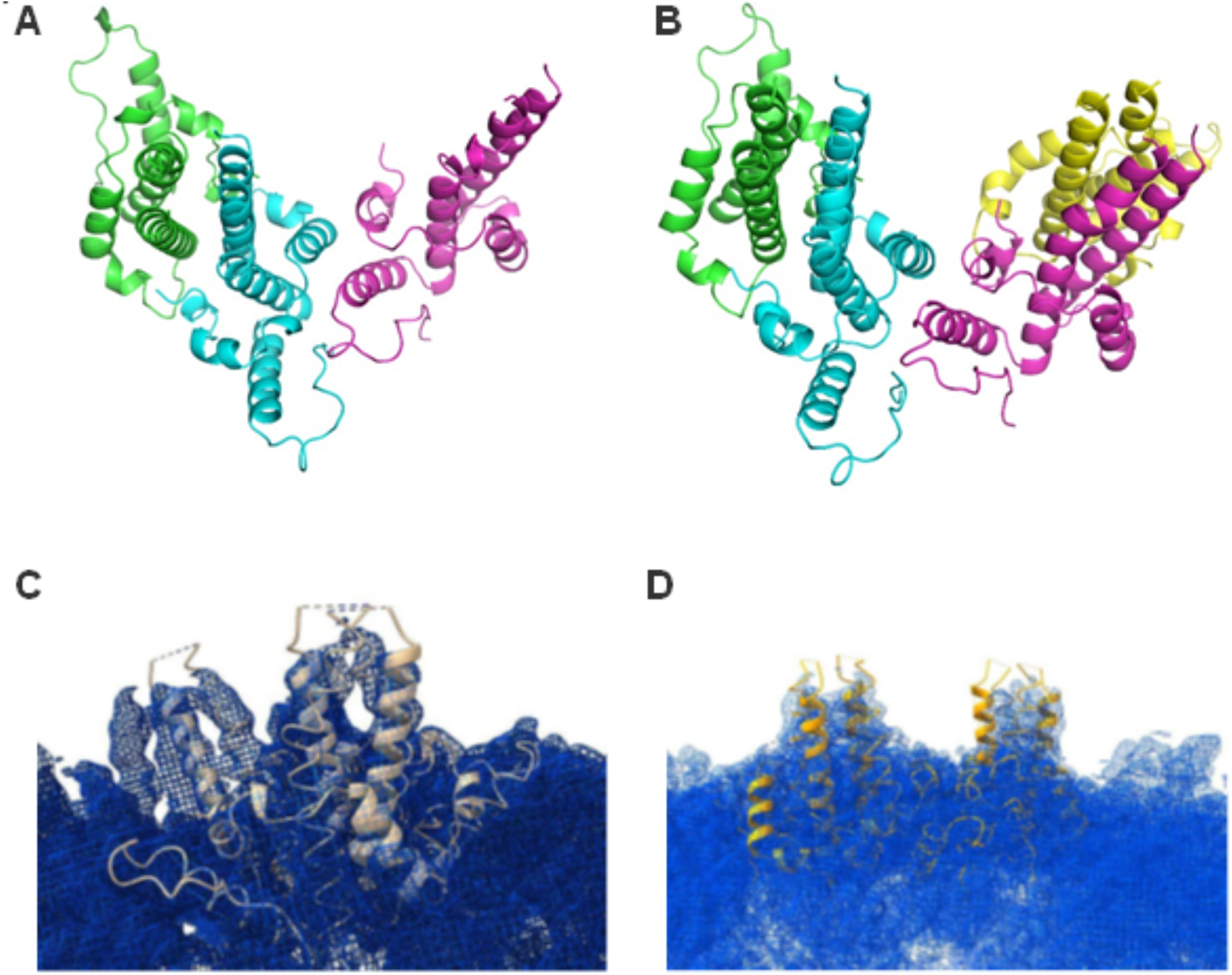
HBc/4M2e monomer structures for T3 (**A**), and T4 (**B**) symmetry and electron density obtained by cryo-EM for particles with T3 (**C**) and T4 (**D**) symmetry inscribed in full-atomic HBc/4M2e model (without residues 75-220).

The study of particle formation using TEM ultrathin sections of bacteria showed that when the HBc/4M2e expressed in the cytoplasm, inclusion bodies are formed, in contrast to HBc without insertion (Figure 6). On the sections of bacteria expressing HBc/4M2e, at the studied expression periods (0, 6, 10, 14h), there is a clear zoning of the bacterial cytoplasm into the central area bounded by a ring of electron-dense material, and rather homogeneous areas of average electron density (they look gray in the photographs), localized, as a rule, on the periphery of the bacterial cell (Figure 6A). The gray homogeneous regions of the cytoplasm shows the typical morphology of inclusion bodies in *E. coli* [12], Thus, according to the data obtained, the predominant part of the overexpressed HBc/4M2e protein apparently aggregates in the form of inclusion bodies. Undisturbed bacterial cells contain inclusion bodies and are morphologically indistinguishable from *E. coli* at earlier stages of overexpression. Ultrastructural analysis of bacteria expressing HBc without insertion revealed that typical inclusion bodies are present in a rather small proportion of bacterial cells (Figure 6B). Most bacteria in their ultrastructure are comparable to bacteria at the “0” point. Thus, when HBc is expressed without a 4M2e insert, most of the overexpressed protein does not form inclusion bodies. Apparently, the assembly of particles took place inside bacteria, immediately after translation, although single spheres are not visible on ultrathin sections.

**Figure 6.**
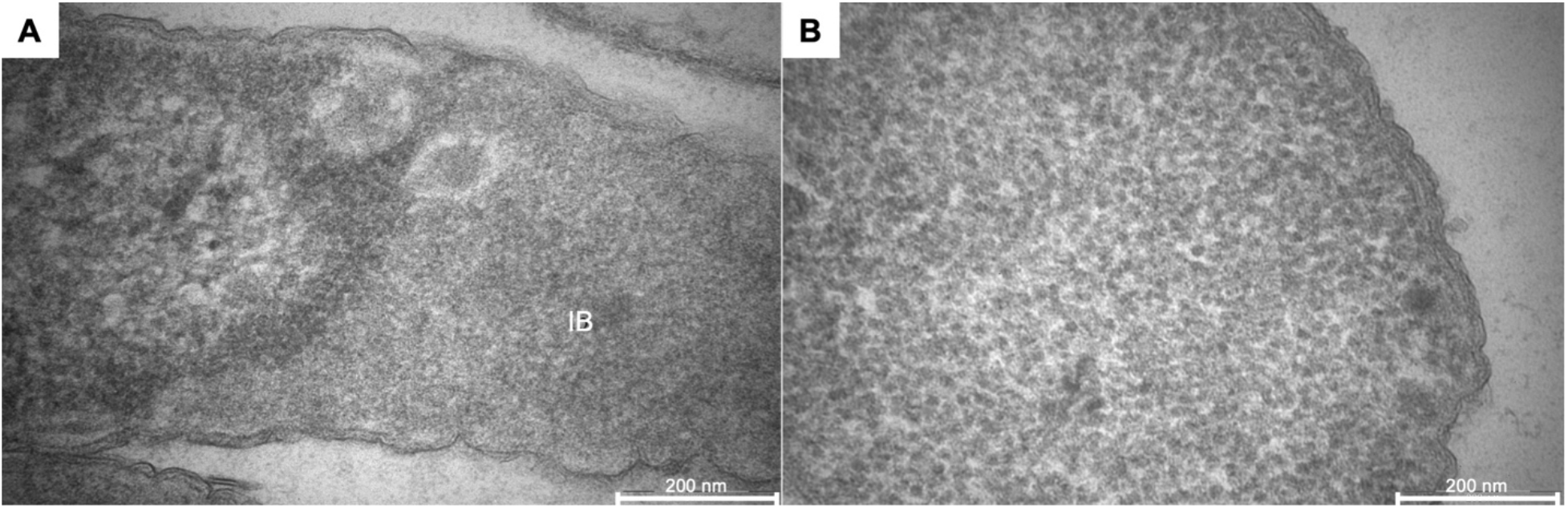
Results of electron microscopy of ultrathin sections of E. coli bacterial cells producing HBc/4M2e (A) and HBc (B) proteins, at 16 hours after expression induction.

All these data indicate the role of the formation of intracellular supramolecular structures in parallel with protein synthesis in the prokaryotic expression system, which raised the question of what is inside the sphere. The control of impurity and the correspondence of the primary protein structure to the proposed was carried out using electrophoretic separation in polyacrylamide gel followed by colloidal Coomassie staining and mass spectrometric identification of the zones corresponding to detected proteins.

The results of PAGE of the recombinant protein HBc/4M2e are shown in Figure 7, the results of mass spectrometry identification of the proteins are summarized in Table 1.

**Figure 7.**
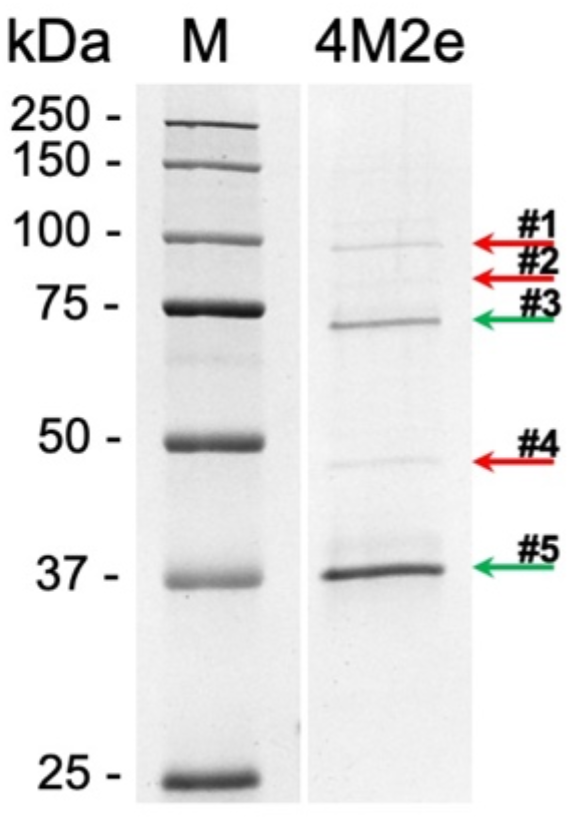
Electrophoretic analysis of protein content in isolated HBc/4M2e particles. Red and green arrows marked bacterial and target protein respectively.

Thus, in the studied fractions, based on mass spectrometry analysis zones corresponding to the electrophoretic mobility of the target recombinant protein (marked with green arrows) were found, and a minor admixture of bacterial proteins (marked with red arrows) in the HBc/4M2e sample was revealed. The content of impurity proteins in the particle did not exceed 6-8%. It should be noted that HBc/4M2e is also presented in dimeric form (arrow #3, Fig.5).

## CONCLUSION

1. M2e antigen used to insert into the HBc antigenic loop, prone to the formation of amyloid-like aggregates.
2. Despite this, HBc/4M2e particles are stable in solution, apparently due to the high surface charge, at least for 1 month.
3. The HBc antigenic loop containing 4M2e does not possess, according to the data of cryo-electron microscopy stable spatial structure. But the 4M2e insertion into the “antigenic loop” is localized outside the HBc/4M2e particle.
4. The HBc/4M2e particles are seem to be synthetized inside the bacterial cytoplasm due to high aggregation propensity of HBc/4M2e monomers. In spite of this, there is no detectable amounts of proteins from the bacterial cytoplasm in isolated HBc/4M2e particle fraction.

## Acknowledgments

The authors acknowledge the support and the use of resources of the Resource Center for Probe and Electron Microscopy at the NRC “Kurchatov Institute”. The study was supported by the Russian Science Foundation project 19-74-20146.

## Author contributions

VVE – writing the manuscript, spectroscopy methods; AVS – cryo-EM data processing; EBP – VLP’s TEM and cryo-EM, data processing; AAS – VLPs purification; YAZ – PAGE, mass-spectrometry; DSV – differential scanning fluorometry; PAN – DLS; ANG – TEM of bacteria; YPG – AFM; AAK – obtaining of bacterial biomass; LAS – peptide design; LMT – project conceptualization and management; TAS – cryo-EM sample preparation; AGM – cryo-EM experiment design and data processing; ALK – project conceptualization and management

## Competing interests

AK is a founder of the company NanoTemper Technologies Rus (St. Petersburg, Russia), which provides services and devices based on MST and nanoDSF and represents NanoTemper Technologies GmbH (Germany). The remaining authors declare that the research was conducted in the absence of any commercial or financial relationships that could be construed as a potential conflict of interest.

